# Concurrent saliency and intentional maps in posterior parietal cortex

**DOI:** 10.1101/2025.11.27.689962

**Authors:** Ruichen Zheng, Yong Gu, He Cui

## Abstract

The goal for motor action represents the desired state of the individual when interacting with the world. Investigating how the goal is generated requires clarifying spatial extrapolation that involves perceptual and action processes. Here, we recorded neural activity from monkeys manually intercepting a circularly moving target being occluded. The saliency and intentional maps, which developed to locate the target of interest and the intended action goal, were thus distinguished in neural activity recorded from the lateral intraparietal cortex (LIP) and parietal reach region (PRR). Dynamic internal representations spanning from visual registration to state estimation were unveiled in both areas, with the goal for the interception being inferred as the visual consequence of the movement. The motion extrapolation was found to be an intrinsic part of movement planning, so that the continuous and stable spatial extrapolation from motion to action was prevalent in both areas. The goal estimation based on the sensory extrapolation rationalized the concurrency and correlation of saliency and intentional maps, which were found to switch their dominance by reorganizing the read-out dimensions of the spatial manifold. The extrapolation achieved at both the perceptual and motor stages not only bridged the saliency and intentional maps, but also revealed the form of the state estimation in guiding predictive sensorimotor control.

## Introduction

Animals live in an ever-changing environment where physical delays in the nervous system cannot be eliminated solely through biophysical improvement. To interact with a dynamic world, the nervous system directs the individual to act toward the future, compensating for delays in the motor stage, as when a predator intercepts the prey (Mischiati, 2015), or regarding primates operating or intercepting a moving object (Flanagan et al., 2001; Zago et al., 2004). Since the flag-lag effect has been observed under various conditions (Nijhawan, 1994; Sheth, Nijhawan, and Shimojo, 2000; Alais and Burr, 2003), whether the motor compensation is sufficient for behavior construction has been considered. As two forms of interaction between the individual and environment, action and perception are argued to be served by separated pathways. If both processes require predictive compensation, identifying the spatial extrapolation they share could be essential in revealing how the motor action is planned.

Controversies surrounding predictive compensation mirror investigations of signals embedded in the posterior parietal cortex (PPC), where extrapolations related to both sensory stimuli and motor consequence have been observed (Assad and Maunsell, 1995; Eskandar and Assad, 1999; Duhamel, Colby, and Goldberg, 1992; Mulliken, Musallam, and Andersen, 2008). As a crucial sensorimotor interface, the PPC has long been suspected to integrate signals from multiple sources to develop a spatial map that locates behavior-relevant motor apparatus and external items, referred to as the intentional map and saliency map, respectively (Andersen and Cui, 2009; Bisley and Goldberg, 2010). The task-dependent literal-to-abstract activity raises rich frameworks centered on action- or perception-oriented processing. However, because the design of conventional tasks caused the motor action to always be directed to predefined sensory cues, the signal reflecting existing stimuli and action goals was thus difficult to distinguish.

In recent studies, a manual interception task was designed to dissociate neural tuning to instantaneous stimulus and upcoming movement (Li, Wang, and Cui, 2018), revealing that pre-movement activity in area 7a co-varies with reach direction instead of target location (Li, Wang, and Cui, 2022). In contrast, motor cortical activity is strongly modulated by target motion (Zhang, Chen, et al., 2024) and is involved in continuous sensorimotor transformations (Zheng, Wang, and Cui, 2025). Nevertheless, it is unclear how predictive signal emerges, and whether it indeed encodes motion extrapolation or action goals. In the present study, we modified the interception task by occluding the target for a fixed period after cue presentation, allowing the movement to be prepared without tracking the target. Our recording from the lateral intraparietal area (LIP) and the parietal reach region (PRR) indicate that interceptive movements were planned in the form of visual extrapolation concerning target motion to movement consequences. The absence of visual input emphasizes the concurrent motion extrapolation and goal estimation, suggesting that sensory extrapolation is integral to action planning. These results suggest the coexistence of saliency and intentional maps, and the transition from the former to the latter, which is achieved by exploiting different dimensions of neural state space. Therefore, seemingly contradictory attention/saliency versus intention/decision models might be reconciled under a continuous sensorimotor transformation, revealing a circuit involving the PPC that converges perception and action stages.

## Results

### Behavior

Two rhesus monkeys were trained to make a reaching movement towards a moving or stationary target while maintaining gaze fixation throughout a trial (Figure 1A). The beginning of a trial was marked by the fixation of gaze and the hand at the center of the screen. After 400 ms, a peripheral target appeared (Cue on) and either moved circularly around the center or remained static. The location of the target was presented during a random Cue period (400-800 ms), and then became invisible (Cue off) during the rest of the trial. After a fixed Occlusion period (500 ms), the fixation dot was dimmed and the monkeys were allowed to reach the peripheral target (Go), and given feedback once a peripheral location was touched.

**Figure 1.**
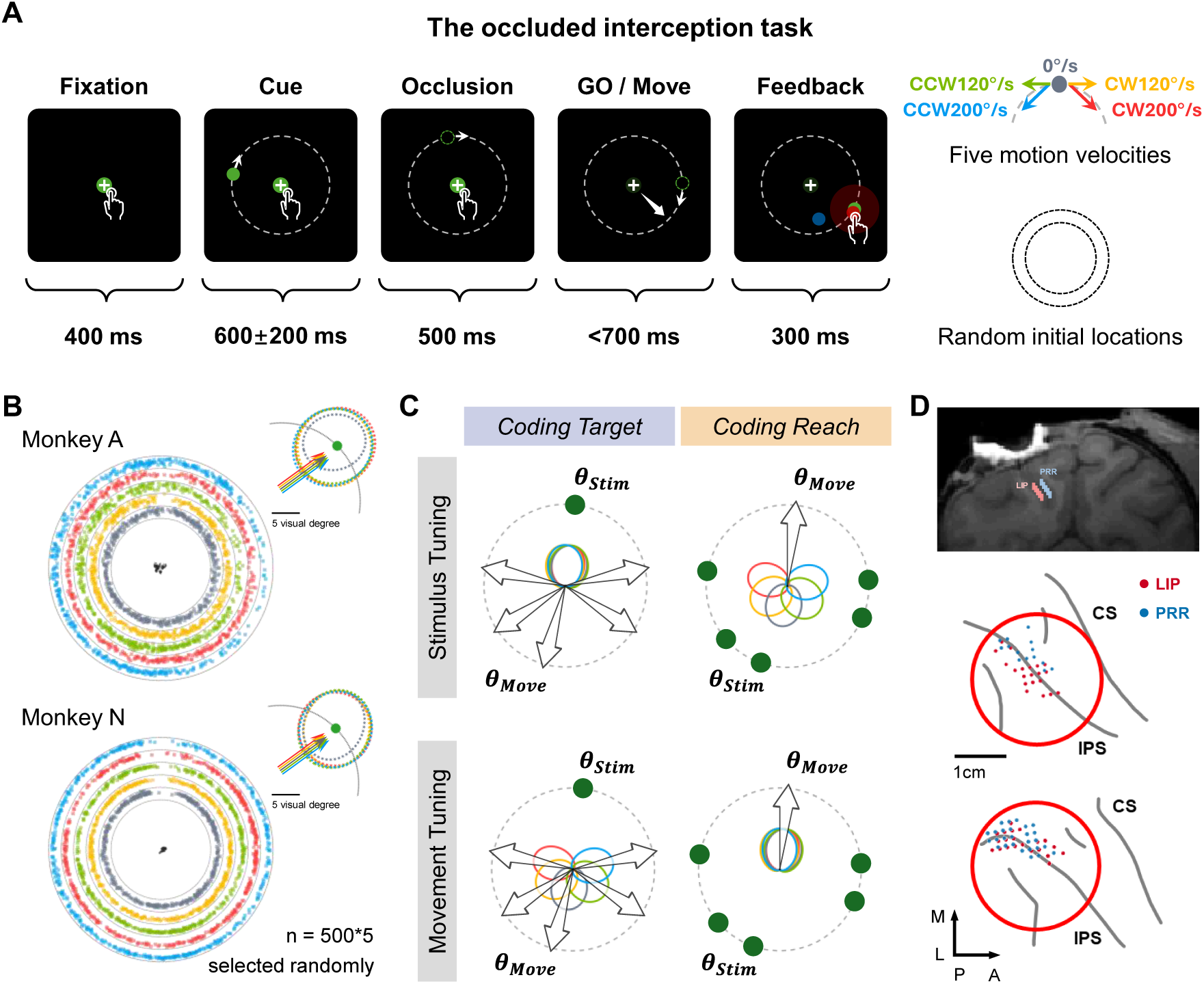
Experimental setup and behavioral performance. (A) The occluded interception task. The monkey faced the visual stimuli displayed on a touch screen and interacted with them by hand. A standard trial consisted of five epochs: Fixation, in which the monkey needed to hold his hand and gaze at the fixation points which denoted by a dot and a cross displaying at the center for 400 ms to start a trial; Cue, during which a target in the form of a green dot appeared at a random location on the peripheral path (Cue on), and the monkey viewed the target, which was either moving or stationary, for 400-800 ms; Occlusion, during which the target disappeared (Cue off), and the monkey waited for another 500 ms for a signal to move; Go/Move, when the central dot was darkened (Go), signaled the monkey to touch the target within 700 ms; Feedback, delivered once the monkey touched the screen for 300 ms, and the location of both the target and touch endpoint were displayed, with the endpoint was colored red and blue for successful and failed trials, respectively. The velocity of the target was randomly selected from 0 °/s, 120 °/s clockwise (CW) or counterclockwise (CCW), and 200 °/s CW or CCW for each trial. The target path with a radius of 14 visual degrees, denoted by the grey dashed circle, was not visible. The central dot and cross comprised fixation points to indicate where the gaze and hand, respectively, needed to be held; The darkness of the dot signified the Go, and gaze was restricted until the end of Feedback. (B) The touch and gaze distribution of monkey A (upper) and monkey N (lower) in successful trials. For visualization, the distribution of reach endpoints under each of the five conditions is scaled and displayed separately in annuli arranged in order, with 500 randomly selected trials for each condition. The annulus indicates a tolerance window of 7 visual degrees, and insets in the top right show the distribution of motor error, which was calculated by aligning the target to the same location (for example, the location marked by the green dot) after the offset between endpoints and the target was acquired. Colored dots and dashed ellipses indicate the endpoints and 95% confidence interval of motor error in corresponding conditions. Gaze trajectories in ten randomly selected trials are denoted by black dots around the center, which wander in the tolerance window of 3 visual degrees. (C) The hypothetical schematic for distinguishing task-related modulation of single units. Stimulus and movement modulation could be identified by comparing the directional tuning curves defined by target direction (upper row) and reach direction (lower row) across five velocity conditions. Since reach endpoints were not coupled with specific target locations, the target passing through the same location could lead to reaching movements in various directions (left column), and movements with the same direction could originate from various target locations (right column). Therefore, a neuron responding to the visual stimulus would show tuning curves directed to the target regardless of the velocity conditions (left column), and a neuron responding to the movement would show tuning curves consistently directed to the reach endpoint (right column). (D) The recording locations. Top panel shows the coronal section that marks the location of the recording chamber. The middle and bottom panels show the lateral view of the right hemisphere of monkey A and monkey N, respectively. Along the intraparietal sulcus (IPS), the estimate of sites for recording LIP and PRR neurons is indicated by red and blue dots, respectively.

The initial direction and the motion velocity of the target were selected pseudo-randomly in each trial, leading to a relatively uniform distribution of reach endpoints (Figure 1B), except that the upper region was relatively less touched due to the experimental apparatus and limb posture. The endpoints did not cluster into certain zones, and were distributed around the target without a significant undershoot or overshoot (the distribution of motor error, i.e., difference between the endpoint and the target position, deviated from zero distribution by less than 1 degree, with no velocity dependence), suggesting that the monkeys did not developed a habit of intercepting only at preferred regions. A difference in eagerness for moving and stationary targets (Figure S1A,B, t-test, p < 0.001) manifested as shorter reaction times and movement duration in the interceptive movement, also implied that the monkeys performed interception with little hesitation.

Intercepting a moving target after significant time without visual information is challenging. After extensive training, the interceptive reach of the monkeys predictively aimed at the target location at the end of the movement, in the absence of habitual movements towards fixed regions, as previously reported in the manual interception of visible targets (Li et al. 2018, 2022; Zhang, Chen et al. 2025). The anticipation required for a successful trial and the flexible relationship between the target and the planned movement make this interception task advantageous for investigating neuronal responses to the visual stimulus and the impending movement.

### Dissociated stimulus and movement modulation in neural responses

Task-related modulation of individual neurons could be described by a tuning curve that specifies the relation between the neuronal activity and the target or movement direction. Benefit from the dynamic visual input and self-initiated movement, organizing the trials according to the instantaneous target direction to fit the directional tuning curves would shuffle the reach direction of corresponding trials, and vice versa. Therefore, at a certain moment, the tuning curves defined by the instantaneous target direction would be aligned across five velocity conditions if a neuron responded to the stimulus, and would be shifted with velocity when defined by the reach direction; conversely, a neuron responding to the movement would demonstrate consistent movement tuning curves and disordered stimulus tuning curves (Figure 1C).

Neurons were recorded from LIP and PRR separately; well-isolated units that displayed task relevance (see Electrophysiology) were included for further analysis (LIP: 86 and 245 for monkey A and monkey N, with single-electrode and linear probes, respectively; PRR: 65 and 294 for monkey A and monkey N, respectively; Figure 1D). In accordance with previous studies involving sensory-guided movement, neurons in LIP and PRR always discharged for a varying period after the stimulus-evoked response. As shown in Figure 2, two example neurons recorded from LIP and PRR, exhibited a rapid elevation in activity following Cue on, and a distinct development of persistent activity as Go approached.

**Figure 2.**
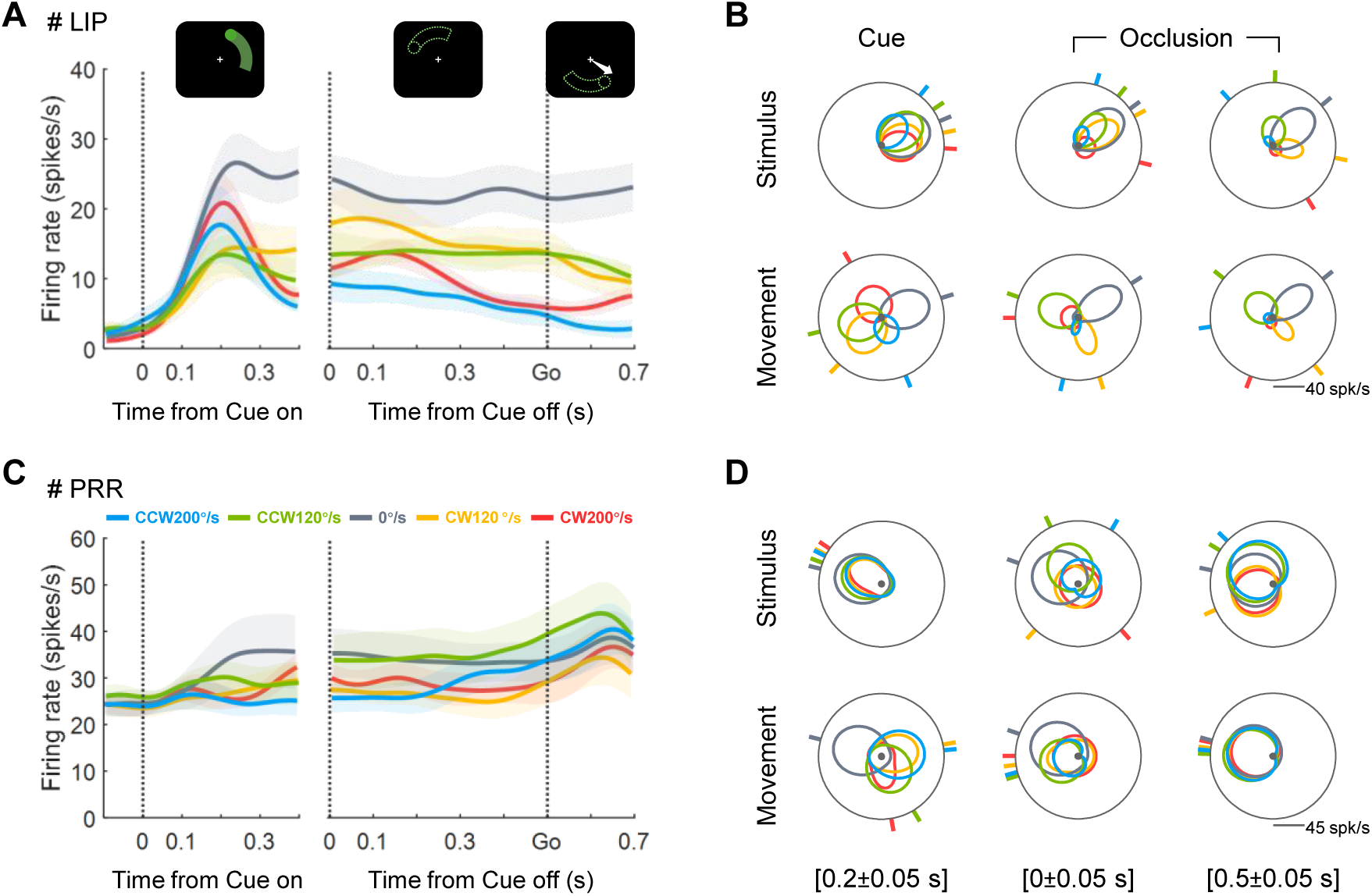
Single-neuron responses during the occluded interception task. (A) Time course of trial-averaged activity for an example LIP neuron. The colored traces represent firing rates averaged across trials in the corresponding conditions. The shaded area indicates the standard error. The vertical dotted lines indicate the time point of three events, i.e., Cue on, Cue off, and Go. (B) The directional tunings of the neuron in (A). The fitted tuning curves and preferred directions (PDs) defined by the direction of the real-time stimulus and reaching movement are shown in the upper and lower rows, respectively. From left to right, the tuning curves under five conditions acquired from time windows centered on 0.2 s after Cue on, Cue off, and Go are denoted by colored petaloid rings, with bars around the circle denoting the corresponding PDs. (C) Time course of trial-averaged activity for an example PRR neuron. (D) Direction selectivity of the neuron in (C).

Activity in both areas appeared to be contingent on the target or intended movement during different periods (Figure S1, S2). Identifying the stimulus- and movement-related signals by directional tuning is consequently essential for revealing the potential representation transition underlying the persistent activity. As exemplified by the two types of tuning around the three task events (Figure 2B and 2D), the vigorous responses triggered by the visual stimulus was reliably tuned to the target location across all conditions, verifying the stimulus modulation at 0.2 s after Cue on in both example neurons. When the target was occluded, the directional tunings diverged between the two neurons. The tuning curves of the LIP neuron were more aligned with the target than the reach direction, while those of the PRR neuron were consistently directed to the reaching movement at both Cue off and Go, indicating an attenuated stimulus modulation and a robust movement modulation in the two neurons, respectively.

### Gradient directional tunings from stimulus to movement

By comparing the stimulus- and movement-defined directional tunings, the time-varying representation preference of each neuron could be assessed. A sensorimotor index, which was defined according to the variance of two types of directional tunings, was computed to quantify the tendency to represent stimulus and movement by a value ranging from -1 to 1. For instance, the index of the two example neurons at 0.2 s after Cue on was -0.9, indicating a strong stimulus modulation; during the Occlusion, the gradually weakened stimulus modulation in the LIP neuron was signified by the index of -0.7 and -0.4, and the constant movement modulation in PRR neuron was signified by the index of 0.8 and 0.9 respectively at Cue off and Go.

From Cue on to Go, the distributions of the index derived from LIP and PRR neurons over different epochs are illustrated in Figure 3A. As shown in six example time windows, the median of the distribution shifted from negative to positive in both populations, suggesting an overall gradient from stimulus coding to movement coding. Additionally, the index of LIP neurons was always significantly lower than that of PRR neurons (time windows labeled with an asterisk, Wilcoxon rank sum test, p < 0.001), indicating that LIP neurons were more likely to be modulated by the stimulus than PRR neurons.

**Figure 3.**
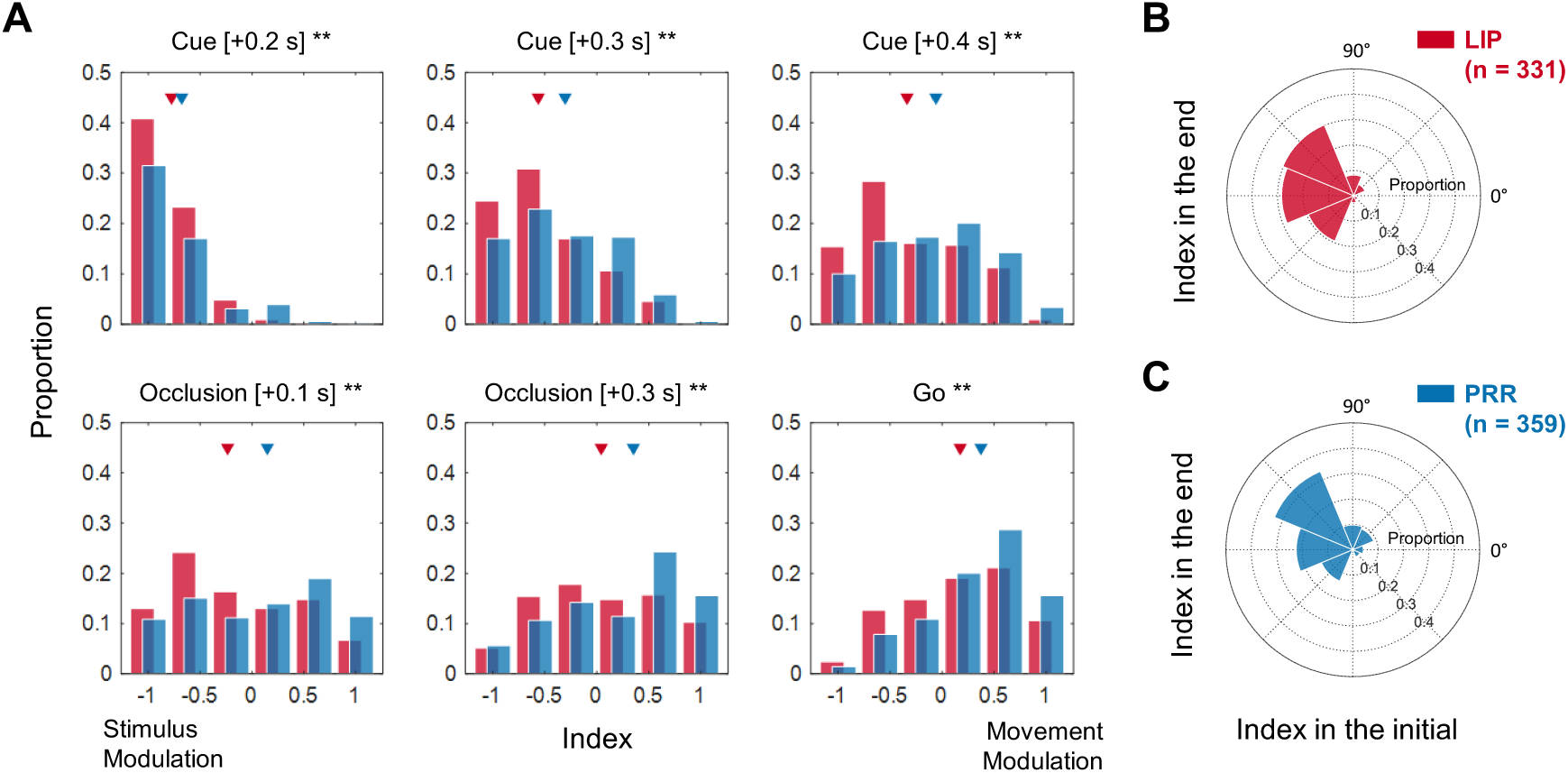
Shift of sensorimotor index in the population of LIP and PRR. (A) Distributions of sensorimotor index across LIP and PRR neurons are represented by red and blue histograms, respectively. In each time bin, the index was counted only when a neuron demonstrated directional selectivity (i.e., direction-sensitive period). The distributions during the Cue, exemplified as time windows centered on 0.2 s, 0.3 s, and 0.4 s after Cue on, are shown in upper panels, from left to right. Distributions during Occlusion, exemplified as time windows centered on 0.1, 0.3 s, and 0.5 s (Go) after Cue off, are shown in lower panels. Triangles indicate the median of the corresponding distributions, and asterisks indicate a significant difference in the index distribution between LIP and PRR neurons (Wilcoxon rank sum test; **, p < 0.01). (B) Comparison of the index at the beginning and end of the direction-sensitive period in LIP neurons. Only neurons demonstrating directional selectivity over 200 ms were counted, whose shift of the index is indicated by a vector determined by the index in the first and last 50 ms of the period. The distribution was calculated by binning vectors into 45° zones. The polar histogram pointing to the angle of 135° indicates the fraction of neurons with an index shift from negative to positive, that is, those neurons displaying sensorimotor transformation. (C) Comparison of the index in PRR neurons.

Does the sensorimotor gradient at the population level result from the separate activation of neurons exclusively responding to the stimulus and the movement? For neurons demonstrating sustained directional selectivity, the shift of the index is denoted as a vector determined by the first and last index in the direction-sensitive period. Distribution of the shift vectors primarily consisted of the sector directing 135°, which includes most of the neurons in two areas, and the sector directing 180°, which is also considerable in the LIP, revealing the subpopulations involved in sensorimotor transformation and sensory processing, respectively (Figure 3B and 3C). Notably, shift vectors are predominantly distributed on the left half of the coordinate axes, implying that the neurons modulated by the movement typically underwent stimulus modulation (i.e., the direction-sensitive period always started from a negative index).

### Courses of tuning state transition

In the tuning gradient from stimulus to movement, some neurons seemed not to be exactly tuned to the instantaneous stimulus or the final movement (for example, the directional tunings in Figure 2B during the Occlusion), raising the question of whether the neurons simply track the actual target when they respond to it.

The neuronal tuning to current target and intended movement was investigated by testing a set of directional variables expressed as the sum of the instantaneous target direction and a deviation angle. The deviation angle is defined by the motion velocity and a condition-independent response latency *τ*, denoting the temporal difference between the represented and the current state of the target. A negative *τ* signifies a neuron tuned to the current target with latency, and a positive *τ* indicates a neuron tuned to the current target in advance. For each time bin, the variables including target states with *τ* ranging from -0.9 s to 0.9 s with steps of 0.05 s and the reach direction were tested to identify those that best explained the directional selectivity of a single neuron. Again, the alignment of the tunings defined by the tested variables across conditions and R-square was used to evaluate the goodness of explanation. The tested variable that maximized the tuning alignment and R-square was defined as the tuning state. Fitting the firing rates by the tuning state significantly improved the R-square (Figure S4, t-test, p < 0.001), confirming that the tuning state was the most probable object to which a neuron was tuned.

Figure 4 shows the tuning states of the example neurons. As suggested by the above comparison of stimulus and movement tunings, the PRR neuron in Figure 2 tracked the target with minimal latency during the Cue, and persistently represented the intended movement during the Figure 4 shows the tuning properties of example neurons. As suggested by the above comparison of stimulus and movement tunings, the PRR neuron in Figure 2 tracked the target with minimal latency during the Cue, and persistently represented the intended movement during the Occlusion (Figure 4B). Such a transition was also observed in LIP neurons (Figure 4C), revealing the predictive compensation concerning both sensory and motor delays at single-neuron level. Other forms of motion extrapolation were also noted; for example, it could be generalized to situations where the visual stimulus was occluded (Figure 4D), accumulated over time (Figure S5A), or maintained stable over-extrapolation (Figure S5B). Delayed visual registration, with or without updating, was kept as well (Figure 4A, S5C). An assortment of tuning state developments suggests a dynamic pool converging salience and intention at different timescales in both LIP and PRR.

**Figure 4.**
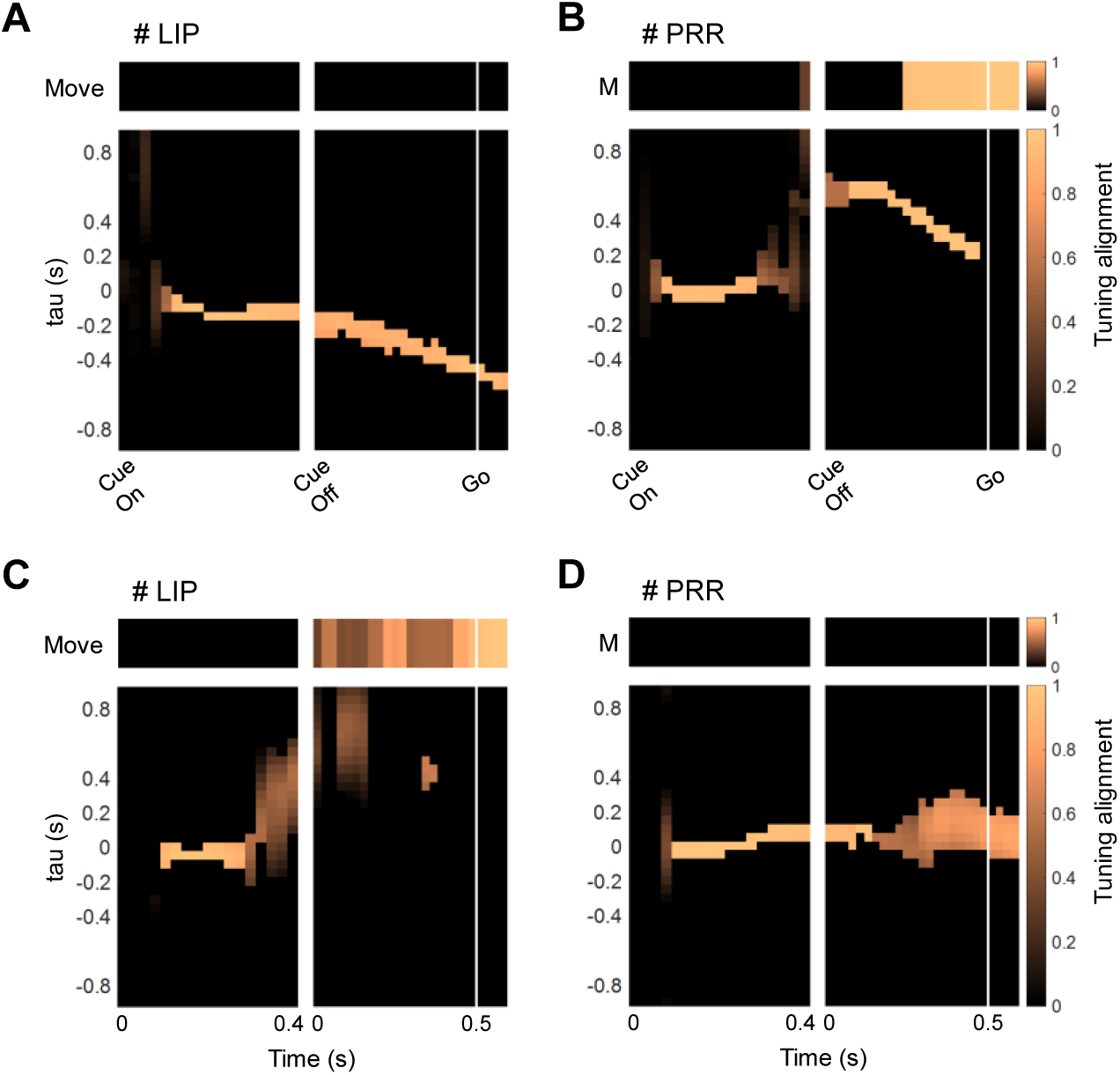
Dynamic tuning states of example neurons. (A) The tuning states of the example neuron in Figure 2B. In each time window, the possibility of a neuron representing the alternative tuning states was indicated by the alignment of corresponding directional tunings between conditions. The higher the tuning alignment, the more likely a neuron is to represent that tuning state, and the brighter the corresponding bin. The alternative tuning states include target states that lead or lag the actual target, signified by “tau” with a positive or negative value, and the target state at the end of the movement, signified by “Move”. For visualization, only tuning states with significantly higher tuning alignment were shown. (B) The tuning states of the example neuron in Figure 2D. (C) The tuning states of another LIP neuron. (D) The tuning states of another PRR neuron.

### Availability of multiple internal estimation in the populations

Since it is not easy to summarize the transition of the tuning state into specific patterns, we first classified the tuning states into four categories according to their temporal relationship with the actual target: the lagging target direction with *τ* less than or equal to -0.1 s; the instantaneous target direction with *τ* between -0.1 and 0.1 s; the leading target direction with *τ* greater than or equal to 0.1 s, but not equivalent to the movement direction; and the intended movement direction. The distributions of tuning states across the two populations are shown in Figure 5A and 5C. In both LIP and PRR, the instantaneous and the lagging states were prevalent during the Cue. As the Cue approached the end, increasingly neurons extrapolated the target states into the near future. After Cue off, the leading state and the movement then became dominant, which was more evident in PRR. The instantaneous motion extrapolation still lasted in the early Occlusion, while the delayed motion information was preserved in a small portion of neurons, which was more common in LIP.

**Figure 5.**
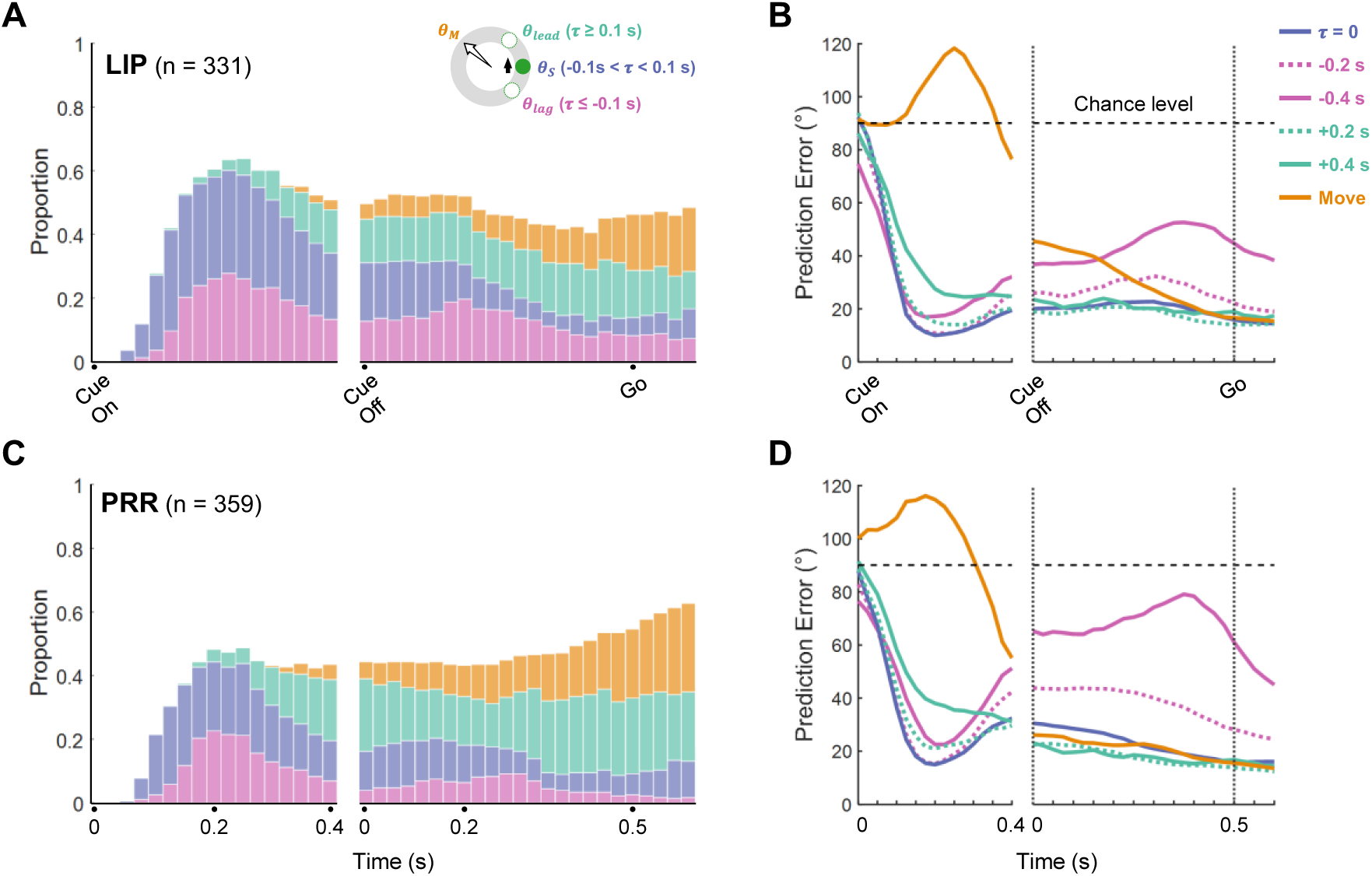
Distribution and read-out of tuning states in the population of LIP and PRR. (A) Proportion of four kinds of tuning state in LIP neurons across time. According to the relation between tuning state and actual target, tuning states related to the target were sorted into lagging, instantaneous, and leading states, depending on whether the τ is larger than 0.1 s or lower than -0.1 s, whose proportions were indicated by histograms colored pink, purple, and cyan, respectively; the proportion of tuning state related to the intended movement is indicated by the orange histograms. (B) Performance of rSVMs derived from LIP neurons during decoding example tuning states. For each time bin, an rSVM was constructed based on the tuning functions of individual neurons. The instantaneous rSVMs were then tested on the neural data to predict alternative tuning states. The error in decoding the instantaneous target direction (τ = 0) and reach direction is indicated by the purple and orange lines, respectively. The error in decoding the direction of lagging states with τ of -0.2 s and -0.4 s is indicated by dotted and solid pink lines, respectively. The error in decoding the direction of leading states with τ of 0.2 s and 0.4 is indicated by dotted and solid cyan lines, respectively. The horizontal dashed line denotes the chance level of decoders. (C) Same as (A), but for PRR neurons. Same as (B), but for PRR neurons.

Next, a set of support vector machine regression models (rSVMs) was constructed to examine the read-out of tuning states from population activity. The rSVM for each time bin was trained by the activity and direction labels interpolated by the instantaneous tuning function of each neuron and tested with real data. The performance for decoding two lagging target states (*τ* = -0.2 s, -0.4 s), the instantaneous target states (*τ* = 0), two leading target states (*τ* = 0.2 s, 0.4 s), and the intended movement was chosen as examples for demonstration (Figure 5B, 5D). Parallel to the distribution of the tuning states, the rSVMs of both areas acquired from the Cue performed well in decoding the location of the instantaneous and lagging target, as well as the leading target. The reach direction was barely embodied in the population activity until the Cue off. During the Occlusion, the accuracy in decoding the movement underwent a gradual and sharp rise in LIP and PRR, respectively, while the accuracy in decoding the instantaneous and leading target remained at a high level. For comparison, decoding the lagging target was no longer as accurate as that during the Cue, especially in PRR.

The tuning states in LIP and PRR demonstrated high flexibility under task restriction, while neurons in LIP registered the target-related salience for a longer period, and neurons in PRR started to figure out the action goal earlier. The availability of visual information facilitated the internal representation to match the actual state of the target; then the visual occlusion prompted the simulation regarding both the motion and movement prevalent across neurons, emphasizing the role of the two areas in the sensorimotor control.

### Reorganization of the potent spatial map underlying sensorimotor transformation

Tuning states changed at heterogeneous pace across single neurons, which benefits selective read-out over a broader range. Considering that the sensorimotor transformation necessarily implicated network computation, the dynamic tuning states could be a manifestation of ongoing computations. However, the directional selectivity, which was determined by the connections, did not undergo significant changes over a short period (Figure S6), raising the question of how the populations modified the internal representation without extensively altering the spatial map.

To further explore the neural population dynamics underlying internal representation of the spatial map, the activity organized by the tuning functions (i.e., tuning matrix) acquired above was transformed into a combination of a set of principal components (PCs) in virtue of the principal component analysis (PCA). As shown in Figure 6A and 6C, the spatial map of the LIP and PRR manifests as a ring consisting of the neural states occupying directions on a circle in the low-dimensional space. As the temporal evolution progressed, the rings colored from blue to green rotated in the state space, demonstrating an invariant spatial map in the first two dimensions and a reorganization in the third dimension.

**Figure 6.**
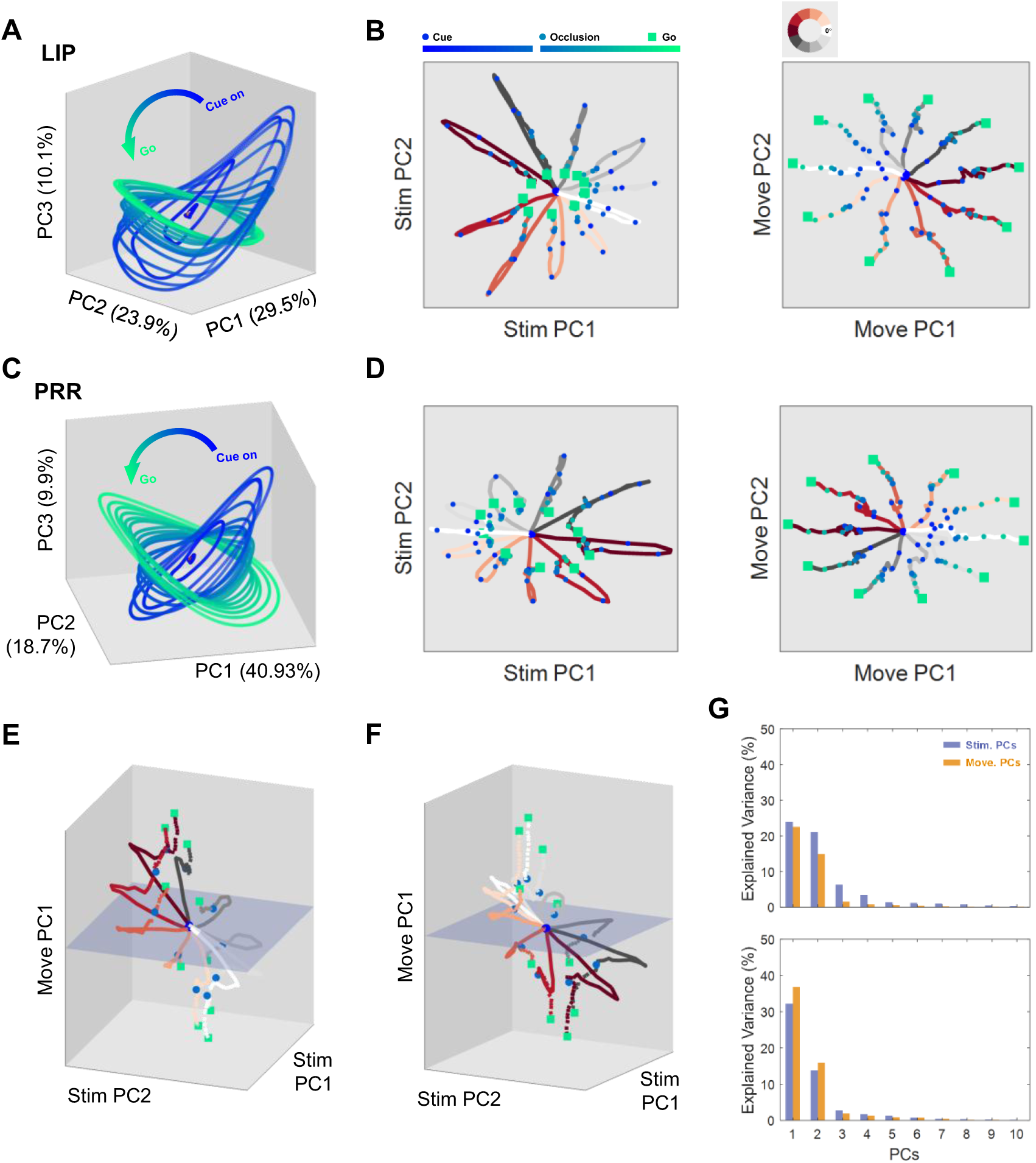
Evolution of the population spatial map in different dimensions. (A) Three-dimensional view of neural states evolution in the tuning subspace of LIP. The ring consisting of neural states distributed in directions from 0° to 360° represents the spatial map of the population in one time section. From Cue on to Go with 50-ms interval, rings colored from blue to green rotated in the low-dimensional space. (B) Projections of the tuning matrix of LIP on the dimensions related to stimulus (left) and movement (right). Traces colored from red to grey denoted the neural trajectory in ten selected directions, with time points marked by dots colored with the same conventions as in (A). (C) Evolution of the spatial map in the tuning subspace of PRR. (D) Projections of the tuning matrix of PRR on the stimulus and movement subspace. (E) Evolution of neural states in the space spanned by the first two stimulus PCs and the first movement PC of LIP. (F) Evolution of neural states in the space spanned by the first two stimulus PCs and the first movement PC of PRR. Proportion of the variance in the tuning matrix captured by the top ten stimulus and movement PCs, which is denoted by purple and orange bars, respectively. The data from the populations of LIP and PRR are shown in upper and lower panels, respectively.

Next, the subspaces related to the stimulus and movement were estimated. One was acquired from the stimulus matrix, encompassing the population activity during the early Cue (from 0.05 to 0.3 s after Cue on) and organized with instantaneous target direction; the other was acquired from the movement matrix, encompassing the population activity during the late Occlusion (from 0.25 s to 0.5 s after Cue off) and organized with reach direction. After performing PCA on the two matrices, the stimulus and movement data were projected onto each resulting subspace. The stimulus PCs captured little variance of the movement-related activity, and the movement PCs captured little variance of the stimulus-related activity (Figure S7, according to Elsayed et al., 2016), suggesting near orthogonality between the stimulus and movement PCs.

With the tuning matrix projected onto the stimulus and movement PCs, we expected that the projections would be active in the dimensions of stimulus subspace during the Cue, and active in the dimensions of movement subspace during the Occlusion. Interestingly, the projections onto both subspaces were active during the Cue. Trajectories in each direction unfolded following Cue on, then folded back in the stimulus subspace and continuously spread in the movement subspace (Figure 6B, 6D). When visualizing the evolution of neural states in the space spanned by the first two stimulus dimensions and the first movement dimension, the transition of the dimensions where the neural states evolved was evident (Figure 6E, 6F).

Together with the similar activity variance explained by the stimulus and movement PCs (Figure 6G), one can speculate that the spatial map locating the behavior-relevant visual stimulus and the motor goal actually overlap, instead of being exclusive in the populations. Switching of the representation could be achieved by exploiting different dimensions of the neural state, which was driven by the temporal dynamics of population activity. Therefore, the neural dynamics related to motion extrapolation and motor planning can be correlated and flexible.

## Discussion

How are perception and action guided under evolving circumstances? In the present study, we designed an occluded manual interception task to prompt the motor action generated in a feedforward manner, disassociating the visual extrapolation and motor planning in the neural representation. A dynamic pool involving visual memory, inferred target motion, and reach planning was observed in both LIP and PRR, with the primary difference between the two areas reflected in the ratio of representations in the populations. The diverse transition from saliency to intentional map was then resolved. Such transitions suggest abundant readout and dynamics in the population activity, which was found to drive the reorganization of the spatial manifold in the neural state space. These findings reveal the mixed spatial maps that located the behavior-relevant sensory cue and motor goal prospectively, demonstrating the engagement of the PPC in a generalized internal model linking perception and action.

Producing proactive motor behavior relies on the estimation of motor consequence, which is known as a forward model (Jordan and Rumelhart, 1992; Wolpert, Ghahramani, and Jordan, 1995), that has been proposed as a solution to overcome the inevitable sensorimotor delay. During interception, the effector could be displaced to the location where the external object will be at the movement offset, demanding simulations for both the object and the effector. The inferred target motion could be conveyed by LIP activity (Assad and Maunsell, 1995; Eskandar and Assad, 1999, 2002), and the modulation of stimulus motion has been observed broadly in retina ganglion cells (Berry et al., 1999), optic tectum (Witten, Bergan and Knudsen, 2006), V1 (Subramaniyan et al., 2018), and V4 (Sundberg, Fallah and Reynolds, 2006). The signal that routes the efference copy of motor command to modulate sensory processing has been observed across species (Crapse and Sommer, 2008), which specifically manifests as remapping of the receptive field (Duhamel, Colby and Goldberg, 1992; Batista et al., 1999), or real-time tracking of an effector (Mulliken, Musallam and Andersen, 2008) in the PPC. However, spatial extrapolation regarding the interaction between object and effector has not been explicitly addressed, although the temporal prediction has been widely investigated under the simplified interceptive movement that the interception zone was pre-determined (Lee, 1976; Port et al., 2001; Merchant, Battaglia-Mayer, and Georgopoulos, 2001, 2003). As in the case of the occluded interception task, a successful interceptive movement was planned based on tracking and predicting the target motion in accordance with limb movement, rendering goal specification to rely on internal prediction instead of artificially defined rules.

The spatial extrapolation concerning target motion and motor consequence was observed in both LIP and PRR. Noticeably, the intended movement was always represented following motion representation, suggesting the correlated extrapolation from motion to movement. The early concept of an internal model emphasized the state estimation in movement generation (Wolpert, Ghahramani, and Jordan, 1995; Wolpert and Flanagan, 2001), that is, the delays in sensory processing and movement execution could be compensated in total by directing action at the desired state (Kerzel and Gegenfurtner_2003; Purushothaman et al., 2008; Sheth and Wu, 2008). In our task, the fixed duration of the Occlusion made the visual extrapolation no longer necessary. Interestingly, visual extrapolation was still observed, while the instantaneous extrapolation that compensated for sensory latency was dominant during the Cue, and the over-extrapolation was dominant during the Occlusion. Although we are unable to determine whether these visual extrapolations are a mix of the delayed target motion and the inaccurate goal estimation, in the view of holistic internal representation, the state estimation at different time-scales, incorporating both motion and motor extrapolation developed in the two areas, suggests a hierarchical predictive coding (Rao, 2024) involving the PPC. The saliency and intentional map have been proposed to be formed in the PPC (Andersen and Buneo, 2002; Bisley and Goldberg, 2010). The intentional map, used to assign the desired endpoint of a motor plan, is inherently anticipatory.

Whereas a large body of evidence suggests that the PPC is composed of multiple functional subareas with distinct roles in the sensorimotor transformation (Kaas and Stepniewska, 2016; Kaas, Qi, and Stepniewska, 2018), we found that LIP and PRR exhibited modest differences in the manual interception task. The LIP and PRR were generally considered to be saccade- and reach-specific, respectively (Snyder et al. 1997; Cui and Andersen 2007). During reach preparation with gaze restriction, a considerable number of LIP neurons were modulated by the forthcoming reaching movement. To examine the potential impacts of the saccadic default plan, the endpoints of the saccades after the trial ended were examined. For both subject monkeys, the directions of the first saccade were rarely correlated with reaching movement (Figure S1C), suggesting that the modulation of intended movement hardly resulted from saccade planning.

A recent study suggests that the LIP and PRR likely plan movements in parallel (Kang, Mooshagian, and Snyder, 2024). In the LIP, reaching movement tuning has been reported in a bimanual movement study (Mooshagiana and Snyder, 2018), and in a task requiring manual operation (Oristaglio et al., 2006; Snyder, Batista, and Andersen, 2000). It is likely that the LIP and PRR plan for the imminent behavior-relevant movement, rather than for the immediately performed movement, which was also the case during decision making, where the LIP activity ramped to form the ultimate decision but not the intervening eye movements (So and Shadlen, 2022). From the viewpoint of the internal model, if the two areas carry the spatial extrapolation pertaining to the effector and target of interest in different modalities, the effector-specific map doesn’t have to be modular, because even movements involving a single effector might cause multisensory consequences (Medendorp and Heed, 2019). Therefore, beyond artificially separating the movement involving different effectors, further studies are required to explore the sensorimotor transformation underlying the ecological behavior.

As discussed above, the saliency and intentional map could be a synthetic spatial manifestation of the behavioral requirement. Considering the abundant connections within the parietal-frontal network, especially the asymmetric projections within the LIP (Ahmed et al., 2025), the saliency and intentional map are expected to be concurrent and intermixed when the PPC fulfills the goal of forward prediction in sensorimotor transformation. In the present study, sensorimotor transformation appeared not to be implemented by sequentially activating separate subgroups solely responsible for sensory processing and movement planning, and the basic direction selectivity did not show significant changes, implying that the saliency and intentional map may share a common spatial map. This could be reasonable at the population level, since sensorimotor transformation needs network computations, while the connections of the networks would not change in a short period. Under the task involving dynamic sensory input and flexible motor output, the sensorimotor transformation was found to be more like a process that defines the spatial map by changing the read-out dimensions. A neuron “possesses” direction selectivity, or the receptive field, before it is specified by task parameters. In the neural state space spanned by the populations, there could be a set of circuits that represent different objects by exploiting different dimensions. The dynamics of population activity was less studied in the PPC. Recently, as the dimensionality of behavioral tasks grows, the dynamics in PPC neurons have also become noticeable (Michaels et al., 2015; Michaels et al., 2018; Diomedi et al., 2021). LIP and PRR neurons exhibited heterogeneity during the occluded interception task, which led to the rotation of the spatial manifold in the state space. The projections of the spatial manifold evolved in both stimulus- and movement-related subspace, suggesting an association between the two kinds of maps. However, the movement information could be read out only during the Occlusion, suggesting a similar mechanism to the null space hypothesis (Kaufman et al., 2014).

As a crucial sensorimotor interface, the converging saliency and intentional maps in the LIP and PRR mediate interactions between individuals and the environment, so the two areas extensively intervene in functions including perception, action, and cognition. In the present study, the goal for motor action was found to be generated based on a sensory extrapolation that could be the foundation of current perception. How spatial extrapolation is determined by task design, how the mental tracking and the motor plan evolve when the monkey views the playback, or performs the interception task with a random Occlusion period, is worth further investigation. Furthermore, to address the utility of the spatial extrapolation, the contribution of these extrapolations to behavioral performance, and how the two areas work together, requires further interventional experiments and large-scale simultaneous recordings.

## Method

### Experimental model and subject details

Two adult male monkeys (*Macaca Mulatta*, 7-9 kg and 9-11 kg for Monkey A and Monkey N, respectively) participated in this study. The monkeys performed tasks in the dark and sat in a consumer-designed primate chair with their heads fixed. A set of titanium (TC4) head post and plastic recording chamber (Crist Instrument, Hagerstown, MD) were implanted stereotaxically by aseptic surgery under isofluorane anesthesia. The locations of the head post and chamber were guided by structural MRI for each monkey. As both monkeys were left-handed, the recording chamber was placed over the intraparietal sulcus (IPS) in the right hemisphere. All training, surgery, and experimental procedures conformed to the National Institutes of Health guidelines and were approved by the Institutional Animal Care and Use Committee (IACUC).

### Task design and apparatus

Behavioral tasks were controlled by a computer running Monkeylogic (http://www.monkeylogic.net) under MatLab (version 2018b, MathWorks). The monkeys viewed and interacted with visual stimuli via a touch screen (EloTouch systems Inc., Menlo Park, CA; 100 Hz sampling rate, 60 Hz refresh rate), which was placed vertically in front of them at a distance of 30-40 cm. Gaze position was monitored by an infrared eye tracker (Eyelink 1000 plus, SR Research, Ontario, Canada; 1000 Hz sampling rate). Neural data was recorded by single tungsten electrode (AlphaOmega) and 64-channel S-probe (Plexon Inc.) for monkey A and monkey N, and collected by AlphaLab SNR (Nazareth, Israel; 22 kHz sampling rate) and Blackrock Microsystems (Cerebus, Salt Lake City, UT; 30 kHz sampling rate), respectively.

In recording sessions, the monkeys performed an occluded manual interception task. To initiate a trial, the monkey was required to foveate and touch fixation points at the center of the screen. After 400 ms, a target was displayed (Cue on) at a random location 14° eccentric from the center, either moving along a circular path with an angular velocity randomly selected from 120 °/s clockwise (CW) and counterclockwise (CCW), 200 °/s CW and CCW, or remaining stationary. The target was visible to the monkey for a random period of 400-800 ms as the Cue, and then became invisible. Once the target was extinguished (Cue off), the monkey kept its hand fixation for another 500 ms as the Occlusion period, and then was instructed to reach the inferred target after the dimming of the fixation point for the hand (Go). A reach (Move) had to be performed within 700 ms, and was considered successful only if the touch endpoint was held within a tolerance window of 7 visual degrees from the target for at least 300 ms. The target stopped once the monkey touched the screen, and reappeared with the touched endpoint as visual feedback after the reaching movement was accomplished (Feedback). The endpoint was replaced by a red dot for a successful reach and a blue dot for a failed reach. The monkey was rewarded with a small amount of liquid food delivered 500 ms after a successful trial. The tolerance windows for fixation of gaze and hand were 3 and 5 visual degree from the center points, respectively, and gaze fixation was restricted until the Feedback ended.

Before or after the recording session, the monkeys were also required to perform mapping tasks consisting of memory-guided delayed saccades and reaches (Snyder et al. 1997). Both tasks were similar to the condition with stationary targets in the interception task, consisting of the Cue period for 500 ms, the Occlusion (Memory) period for 900 ms, the Move period for 700 ms, and the Feedback period for 300 ms. The mapping tasks required the monkey to make eye movement or arm movement to the memorized target according to the color of the target and fixation point, with yellow and green indicating the saccade and the reach, respectively. Gaze was restricted in the reach task, as in the interception task, and the hands were restricted in the primate chair in the saccade task.

### Electrophysiology

The activity of well-isolated single units in the lateral intraparietal area (LIP) and parietal reach region (PRR) was recorded using single-channel electrodes and multi-channel linear electrodes for monkey A and monkey N, respectively. Total of 97 recording sessions were conducted in monkey A and 120 in monkey N. In each recording session, an electrode was directed through a grid to target the lateral or medial bank of the IPS using a motorized microdrive (Thomas Recording GmbH MEM, Giessen, Germany). The subarea to which each recorded neuron belonged was identified in combination with anatomical location and physiological criteria (Andersen, Martyn Bracewell, et al., 1990; Snyder, Batista, and Andersen, 1997). The putative LIP and PRR sites were roughly distributed 3.5-8 mm from subsurface, 2.3–13 mm from midline to lateral, and -10 mm (posterior) to +3.5 mm (anterior) relative to the interaural plane. Neurons with spatial selectivity that discharged persistently during the mapping tasks were further characterized as LIP or PRR according to the effector preference, and were included in the dataset.

### Analysis of behavioral data

Monocular position signals were recorded in real-time and were analyzed off-line. The gaze signal was first filtered with a low-pass Butterworth filter to remove the high-frequency noise above 40 Hz. The filtered signal was then smoothed with a 15-ms width rectangular filter. For each trial, the signal in the period from Feedback to Reward was selected to inspect the saccades.

After the Feedback, the saccades were detected by the threshold of 30 to 400 visual degrees per second for the horizontal or vertical component. Saccades with a duration of less than 15 ms were excluded, as were saccades with an amplitude greater than 40 (Li, Wang, and Cui, 2018).

### Single-neuron analysis

Single units were isolated from the filtered neural signal using Spike2 (v7.15, Cambridge Electronic) for data collected by single-electrode recordings and Kilosort 3.0 (Pachitariu et al., 2016) for data collected by multi-channel recordings.

After acquiring the spikes for each unit, the 1 ms-discretized spike train was aligned to Cue on and Cue off, and was convolved with a Gaussian kernel (δ = 50 ms) to generate peri-stimulus time histograms (PSTHs). Trial-averaged PSTHs were acquired by averaging the activity from trials grouped by velocity conditions or direction zones. The standard error was calculated using 10 bootstraps sampled from the corresponding group of trials.

For further analysis, the firing rates were calculated for each trial in a 50-ms time window sliding with 25-ms steps, and were then standardized by Z-score to ensure that the effect of overall activity fluctuation would not dominate subsequent analyses. After simplifying the visual coordinate to the direction variable, the directional tuning curves were obtained by fitting the firing rates with the direction variables on corresponding trials by the circular version of the Gaussian function:

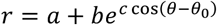

where *r* is the firing rate in a given time window; *a*, *b*, *c* are coefficients; *θ* is the direction of the interested variable, which can be the instantaneous target direction or the reach direction, and *θ*_0_ is the direction that maximizes the firing rate, i.e. the preferred direction (PD) defined by the interested variable. Directional tuning curves were fitted for each condition, and were filtered by the goodness-of-fit (R-square > 0.1). Given that direction selectivity is considered invariant across conditions, the alignment of directional tunings, as measured by the variance of the PDs or the averaged correlation between the tuning curves, could be used to assess the response preference of a neuron. The preference to respond to the stimulus or the movement was measured by the sensorimotor index, defined as follows:

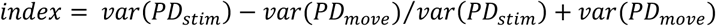

where the *PD*_*stim*_ and *PD*_*move*_ are PDs defined by the instantaneous target direction and the reach direction, respectively. Indexes close to -1 and 1 indicate that a neuron selectively responds to the stimulus and the movement, respectively. The index was calculated only when a neuron showed directional selectivity (the neural responses significantly varied with directions: Kruskal-Wallis test, p < 0.01; R-square of the fitted directional tuning > 0.1). For the neurons displayed directional selectivity for at least 200 ms, the index shift was indicated by a vector whose coordinate was the index at the initial and the last 50 ms of the direction-sensitive period.

The variable that modulates the neural activity may not be confined to the current target or the upcoming movement. In this instance, the variable of interest, *θ*, could be replaced by *θ*_*s*_ + *τν*, where *θ*_*s*_ is the instantaneous target direction, *ν* is the motion velocity, and *τ* is the condition-independent response latency. The target directions with *τ* ranging from -0.9 s to 0.9 s with 0.05 s step, as well as the reach direction, were tested to find the variable that was most likely to modulate the neural activity. The directional tunings corresponding to the alternative variables were fitted within and between conditions. The correlation between the tunings fitted under individual conditions, together with the R-square of the tuning fitted across conditions, was used to evaluate the tested variables. The direction variable that maximizes the tuning correlation and R-square was defined as the tuning state, and the corresponding function was defined as the tuning function.

### Population Analysis

To enable the activity of single units to be processed as the population activity recorded simultaneously, the normalized firing rates were interpolated through the tuning function of individual neurons. The interpolated data from each area was then integrated to constitute a tuning matrix with N, the number of neurons, C, the number of conditions (usually the directions), and T, the number of time points.

The readout of alternative tuning states from the population activity was examined by support vector machine regression models (rSVMs) constructed based on the instantaneous tuning functions of each neuron. The training set was the tuning matrix arranged by the direction of tuning states. The direction labels, ranging from 0° to 360° with 10° interval, were transformed into a combination of the horizontal component *x* = *radius* ∗ cos (*θ*) and the vertical component *y* = *radius* ∗ *sin*(*θ*), where the target path defined the radius. Therefore, two independent decoders corresponding to each linear component were trained, and then tested with real data. The predicted values were converted into directions by the anti-trigonometric function, and the absolute difference between the predicted and the true values was defined as the prediction error to indicate the decoding performance. A permutation test was performed by randomly shuffling the direction labels across trials and recalculating the prediction error 1000 times to acquire the chance level of the rSVMs.

The latent factors in population activity were revealed by transforming the tuning matrix into a set of principal components (PCs) through the standard principal component analysis (PCA).

Before the dimension reduction, the mean activity in the dimension of time was subtracted, and the dimensions of condition were arranged according to tuning directions. The resulting neural states at a given moment were referred to as the spatial map of the population.

The typical task-related dimensions were identified through two other activity matrices which were related to the stimulus and movement, respectively. The former segmented from the population activity during the early 250 ms of the Cue (from 50 ms to 300 ms after Cue on), organized according to instantaneous target direction; the latter segmented from the population activity during the late 250 ms of the Occlusion (from 250 ms to 500 ms after Cue off), organized according to reach direction. The PCs acquired by applying PCA on each matrix were referred to as stimulus and movement PCs, respectively.

For all three subspaces, accumulated variance explained by PCs exceeded 85% of the total by the tenth PC; therefore, the tuning, stimulus, and movement subspaces were defined by the top ten PCs of each. The orthogonality of the stimulus and movement subspaces was quantified according to the variance shared between the two (Elsayed et al., 2016), and the dynamics of the latent factors of population tunings on task-related dimensions was explored by projecting the tuning matrix onto the movement- and stimulus-PCs.

---

**Figure S1.**
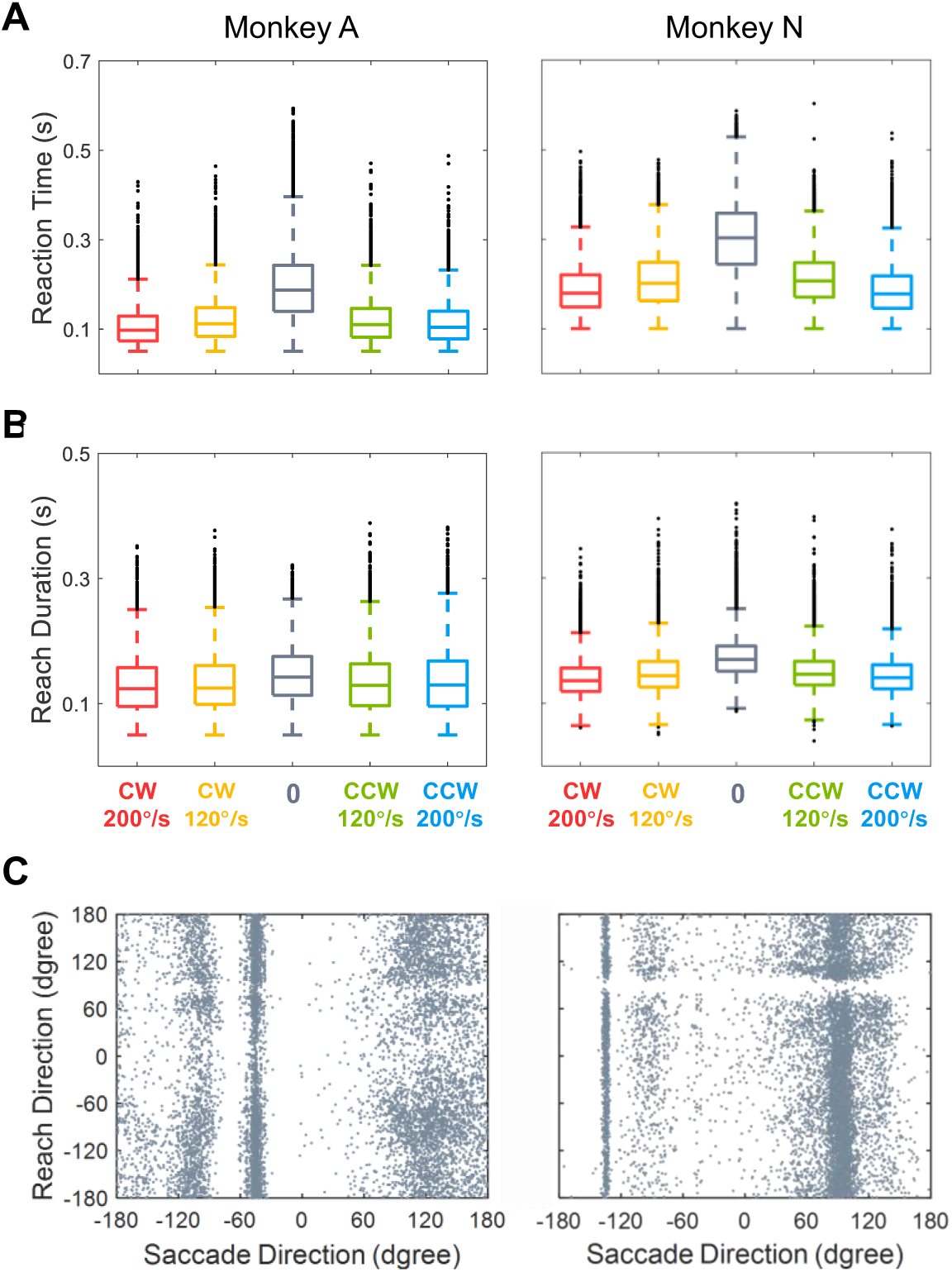
Reaching and saccadic behavioral responses. (A) Reaction time of monkey A (left) and monkey N (right) in successful trials. From left to right, the distribution in conditions of 200 °/s and 120 °/s CW, 0 °/s, 120 °/s and 200 °/s CCW was shown by the colored box. (B) Reach duration of monkey A (left) and monkey N (right) in successful trials. (C) Relationship between direction of the first saccade after trial end and reaching movement. For this demonstration, data were selected randomly from all successful trials in recording sessions, with 2000 trials for each condition. The correlation between saccadic and reaching directions was negligible (linear regression, R-square < 0.01, p > 0.1).

**Figure S2.**
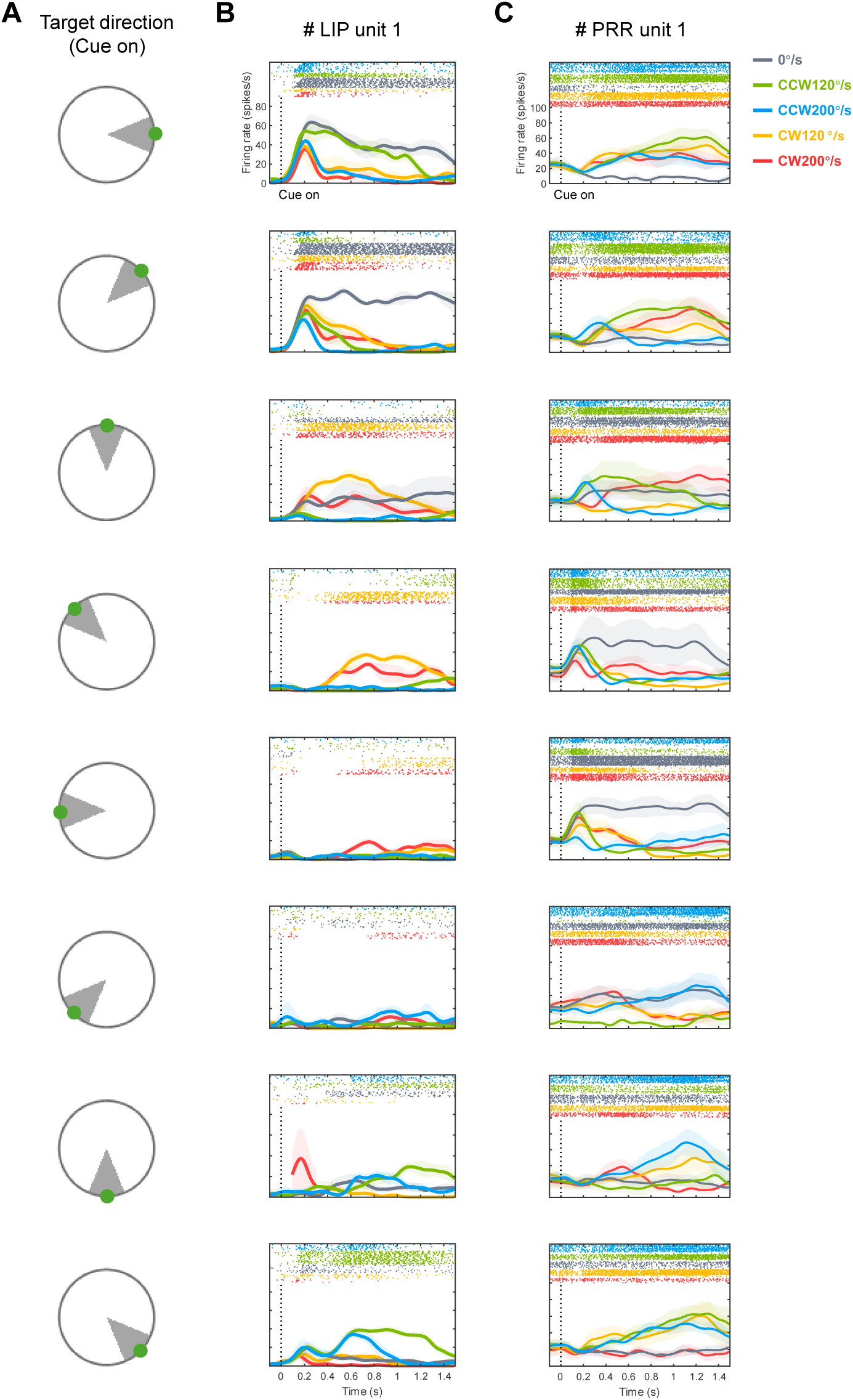
Trial-averaged responses of the neurons in Figure 2 under different velocity conditions and initial regions of the target. (A) Schematic of direction grouping. From 0° to 360°, directions were zoned into eight groups at 45° interval according to the initial target location. (B) Activity of the neuron in Figure 2A when aligning spike trains to Cue on. The panel in each row displays the neuronal responses in the condition in which the target moved from the zone indicated by (A), with velocity indicated by color. (C) Same as (B), but for the neuron in Figure 2B.

**Figure S3.**
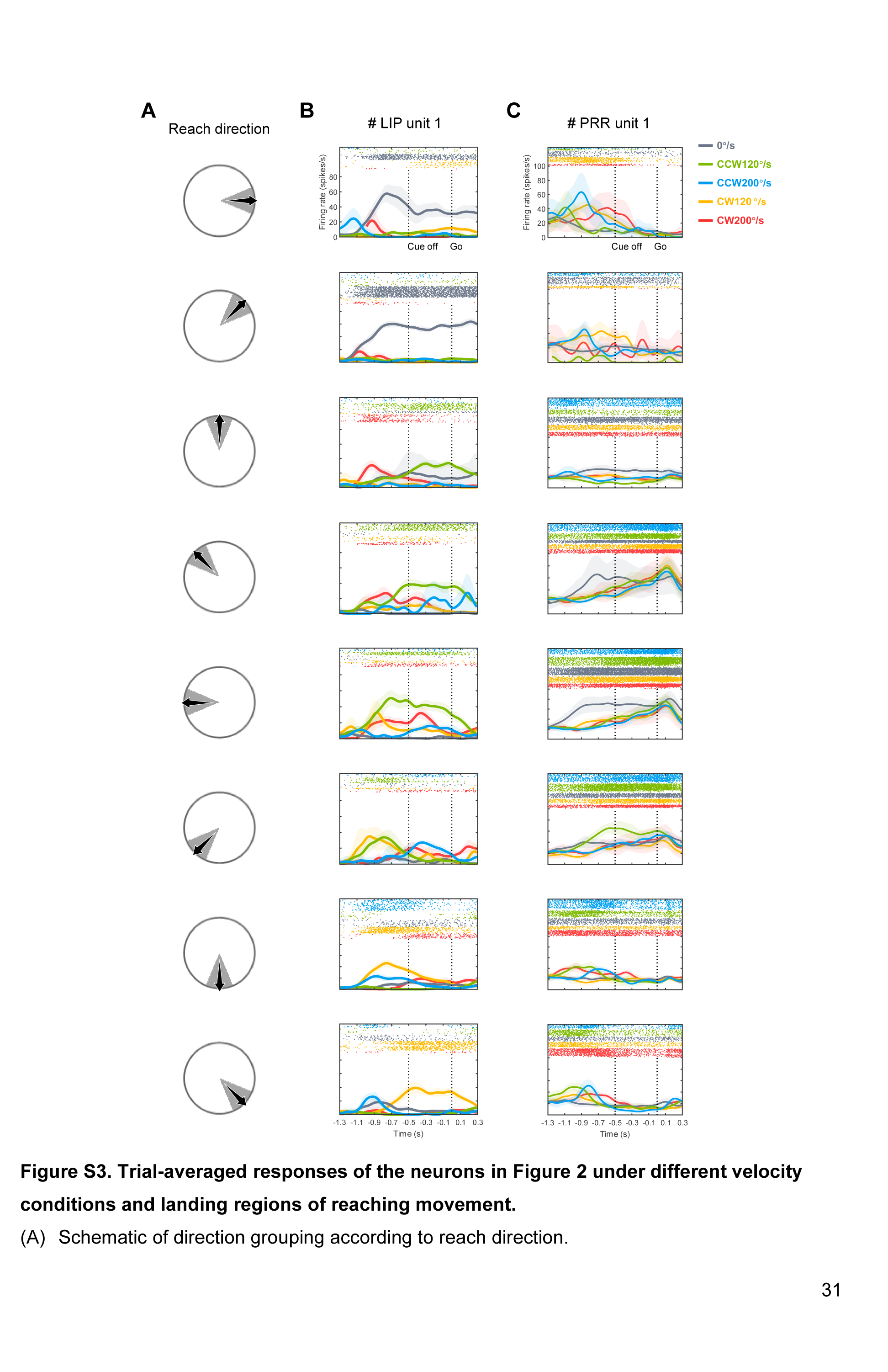
Trial-averaged responses of the neurons in Figure 2 under different velocity conditions and landing regions of reaching movement. (A) Schematic of direction grouping according to reach direction. (B) Activity of the neuron in Figure 2A when aligning spike trains to Go. The panel in each row displays the neuronal responses in the condition that the reach landing in the zone indicated by (A) to intercept the target moving with velocity indicated by color. (C) Same as (B), but for the neuron in Figure 2B.

**Figure S4.**
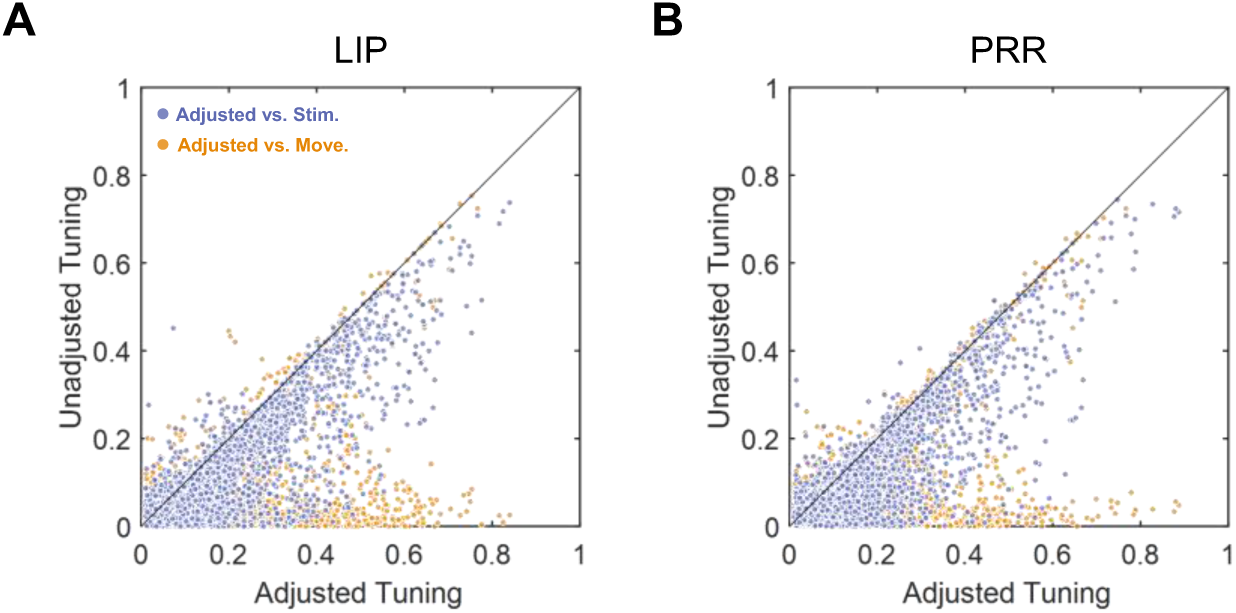
Goodness-of-fit before and after fitting the firing rates with tuning states. (A) R-square of directional tunings of LIP neurons, calculated without grouping trials by conditions. Purple dots represent the comparison between tunings fitted by the instantaneous target and tuning state, with the R-square of each pair plotted as ordinate and abscissa, respectively. Orange dots represent the comparison between tunings fitted by the movement and tuning state. (B) Same as (A), but for PRR neurons.

**Figure S5.**
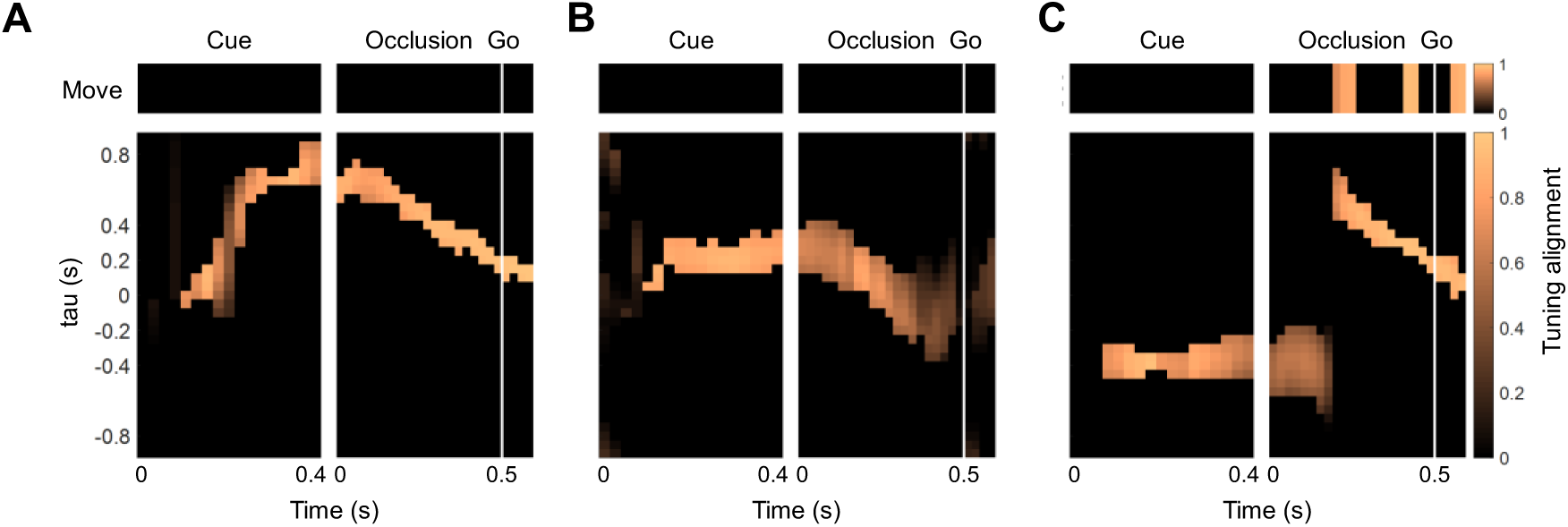
Dynamic tuning states of other neurons. Same as Figure 5, but for neurons in (A) monkey N, PRR; (B) monkey A, PRR; (C) monkey A, PRR.

**Figure S6.**
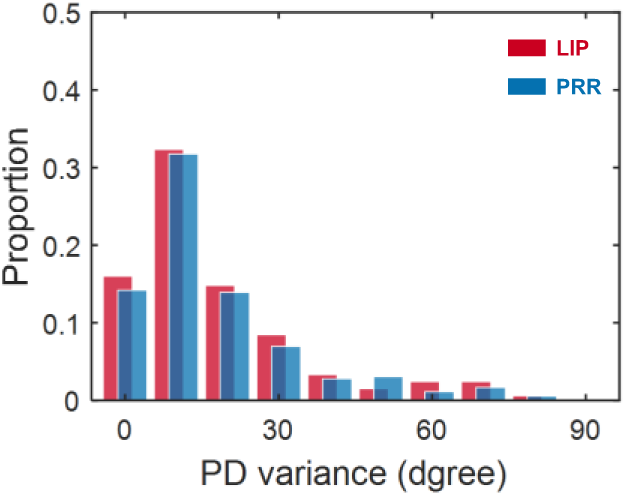
Variance of single neurons’ directional selectivity. Distribution of the variance for preferred direction in the direction-sensitive period. Red and blue histograms represent variance distributions in LIP and PRR, respectively. Variance was calculated only when a neuron showed directional selectivity over 200 ms.

**Figure S7.**
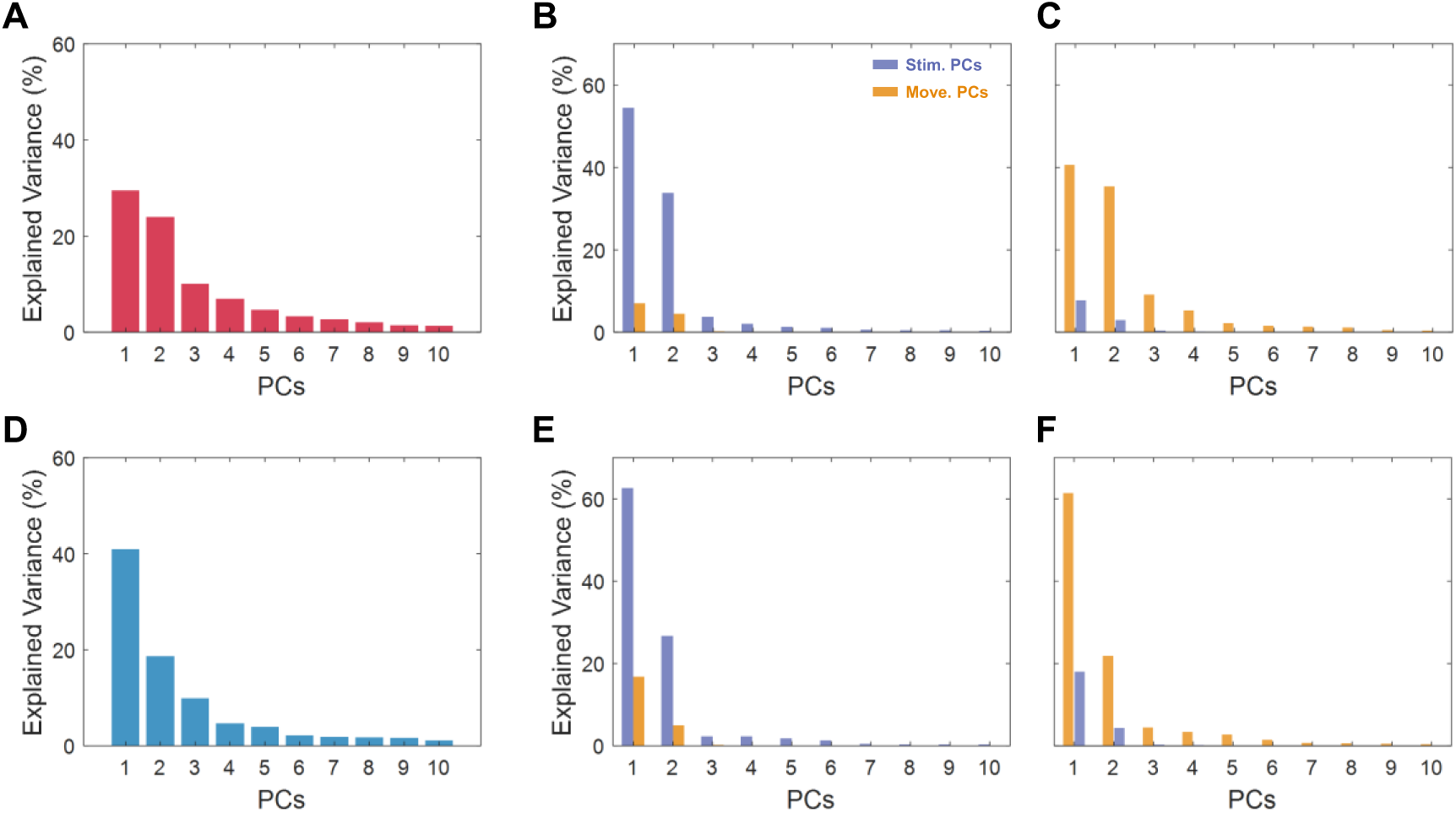
Activity variance explained by the principal components of different subspaces. (A) Fraction of variance explained by the top ten principal components defined by population tuning data. (B) For the stimulus-related data, the fraction of variance captured by the top ten principal components of stimulus and movement subspace (Stim-PCs and Move-PCs) is indicated by the purple and orange bars, respectively. (C) For the movement-related data, the fraction of variance captured by the top ten stimulus and movement principal components. (D) Same as (A), but for the population activity of PRR. (E) Same as (B), but for the population activity of PRR. (F) Same as (C), but for the population activity of PRR.

**Table S1.** Statistical data of sensorimotor index distribution under six time windows.

|  |  | Cue +0.2 s | Cue +0.3 s | Cue +0.4 s | Occlusion +0.1 s | Occlusion +0.3 s | Occlusion +0.5 s / Go |
| --- | --- | --- | --- | --- | --- | --- | --- |
| LIP | Skewness | 1.45 | 0.68 | 0.34 | 0.29 | 0 | -0.20 |
|  | Difference from zero | p<0.001 | p<0.001 | p<0.001 | p<0.001 | p=0.07 | p<0.001 |
| PRR | Skewness | 1.24 | 0.27 | -0.03 | -0.18 | -0.46 | -0.60 |
|  | Difference from zero | p<0.001 | p<0.001 | p=0.01 | p=0.05 | p<0.001 | p<0.001 |
| Difference between areas |  | p<0.001 | p<0.001 | p<0.001 | p<0.001 | p<0.001 | p<0.001 |

## Notes

### Competing Interest Statement

The authors have declared no competing interest.

### Summary of Updates

Modified some wording in Results. Figure legends revised; Discussion updated.

### Funding

- National Science Foundation of China: T2596043, 31671075

